# A covariation analysis reveals elements of selectivity in quorum sensing systems

**DOI:** 10.1101/2021.03.25.437039

**Authors:** S Wellington Miranda, Q Cong, AL Schaefer, EK MacLeod, A Zimenko, D Baker, EP Greenberg

## Abstract

Many bacteria communicate with kin and coordinate group behaviors through a form of cell-cell signaling called acyl-homoserine lactone (AHL) quorum sensing (QS). In these systems, a signal synthase produces an AHL to which its paired receptor selectively responds. Selectivity is fundamental to cell signaling. Despite its importance, it has been challenging to determine how this selectivity is achieved and how AHL QS systems evolve and diversify. We hypothesized that we could use covariation within the protein sequences of AHL synthases and receptors to identify selectivity residues. We began by identifying about 6,000 unique synthase-receptor pairs. We then used the protein sequences of these pairs to identify covariation patterns and mapped the patterns onto the LasI/R system from *Pseudomonas aeruginosa* PAO1. The covarying residues in both proteins cluster around the ligand binding sites. We demonstrate that these residues are involved in system selectivity toward the cognate signal and go on to engineer the Las system to both produce and respond to an alternate AHL signal. We have thus demonstrated a new application for covariation methods and have deepened our understanding of how communication systems evolve and diversify.

---

Quorum sensing (QS) is a widespread form of cell-cell signaling that bacteria use to coordinate the production of public goods including toxins, antibiotics, bioluminescence, and secreted enzymes (Waters and Bassler, 2005; Whiteley et al., 2017). Many Proteobacteria (Case et al., 2008) and Nitrospirae (Mellbye et al., 2017) employ a form of QS based on acyl-homoserine lactone (AHL) signals. AHL QS systems consist of two proteins: a LuxI-type signal synthase and a LuxR-type receptor (Figure 1a). The signal synthase produces an AHL from S-adenosylmethionine (SAM) and an acyl-acyl carrier protein (ACP) for some LuxI-type synthases or an acyl-coenzyme A (CoA) substrate for others (Schaefer et al., 2008) (Fig. 1b). AHL signals can freely diffuse through cell membranes (Kaplan and Greenberg, 1985; Pearson et al., 1999) and at low cell density the QS system is “off”. At high cell density, the signal accumulates and binds the LuxR-type receptor which is a cytosolic transcription factor that regulates gene expression in response to signal binding.

**Fig. 1.**
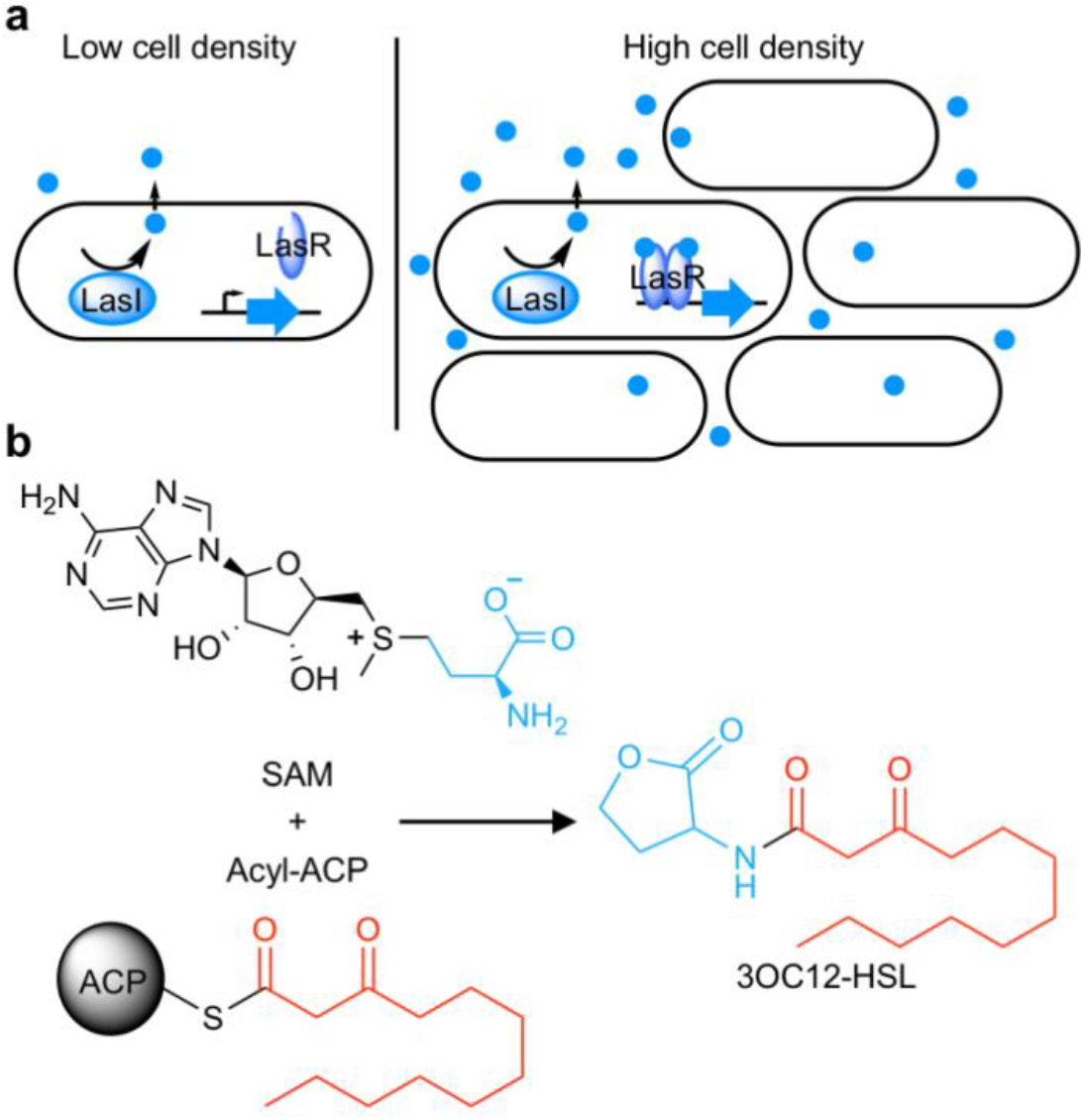
Schematic of acyl-homoserine lactone (AHL) quorum sensing (QS). **a)** AHL QS circuits consist of a signal synthase, a LuxI homolog (LasI in this cartoon), that produces an AHL signal. At low cell densities, the system exists in an “off” state. At high cell densities, the AHL concentration increases and the signal binds its cytosolic receptor, a LuxR homolog (LasR in this cartoon), which functions as a transcription factor. **b)** LuxI-type synthases produce AHL signals from two substrates: S-adenosyl methionine (SAM) and an activated organic acid in the form of an acyl-acyl carrier protein (ACP) or acyl-coenzyme A (CoA). SAM provides the lactone core, which is conserved across all AHL signals, while the acyl-ACP (shown here) provides an acyl moiety which varies between signaling systems. In this example, the synthase LasI produces *N*-3-oxo-dodecanoyl-L-homoserine lactone (3OC12-HSL).

AHL signals share a conserved lactone core, but vary in the acyl moiety which can be a fatty acid ranging from 4 to 20 carbons long, with potential oxidation on the C3 carbon and varying degrees of unsaturation, or can have an aromatic or branched structure (Rajput et al., 2016). This variability in the acyl portion of the signal confers selectivity to the system. Typically, a LuxI-type synthase produces a primary AHL to which its paired LuxR-type receptor selectively responds (Aframian and Eldar, 2020). Selectivity is critical to cell signaling in order to avoid undesired cross-talk or spurious outputs (Laub, 2016). In the case of QS, selectivity ensures bacteria cooperate only with kin cells.

Despite its importance, we know little about how QS systems achieve selectivity or how they evolve and diversify to use new signals. Although the conserved amino acids essential for synthase and receptor activity are well described (Parsek et al., 1997; Zhang et al., 2002), residues that dictate selectivity are often different from those that are required for activity (Collins et al., 2005). Due to the low amino acid sequence identity between LuxI/R homologues, it has been difficult to determine how QS systems discriminate between various AHL signals (Fuqua et al., 1996).

We hypothesized that we could use covariation patterns to identify QS selectivity residues. Such methods have been used to identify amino acid residues that interact with each other within proteins and between proteins that physically bind each other (Aakre et al., 2015; Ovchinnikov et al., 2014; Skerker et al., 2008). Here, we endeavored to expand these methods to assess the interaction between AHL synthases and receptors. While AHL synthases and receptors do not physically interact, they interact indirectly via binding to a shared cognate signal and are believed to coevolve to maintain this shared signal recognition (Aframian and Eldar, 2020). Phylogenetic analyses also support coevolution of synthases and receptors (Gray and Garey, 2001; Lerat and Moran, 2004). We therefore hypothesized that we could identify amino acid residues that covary between QS synthases and receptors, and further, that the covarying residues would be those responsible for signal selectivity.

We used a statistical method, GREMLIN (Kamisetty et al., 2013), to measure covariation within the sequences of AHL synthase-receptor pairs and mapped the covarying residues onto the LasI/R QS system of *Pseudomonas aeruginosa* PAO1. By mutating the top-scoring residues identified by GREMLIN, we demonstrate that they are indeed important for signal selectivity and, further, that these residues can be used to rationally engineer LasI/R to produce and respond to a non-native signal. We thus demonstrate a new application for powerful covariation methods and at the same time identify determinants of QS selectivity.

## Results

### Covariation patterns in QS systems

To begin our analysis, we gathered select protein sequences for known synthase-receptor pairs (Supplementary Table 1) and used these sequences to search the European Nucleotide Archive (ENA) database (Amid et al., 2019) from the European Bioinformatics Institute and the Integrated Microbial Genomes and Microbiomes (IMG/M) database (Chen et al., 2020) from the Joint Genome Institute (JGI) for additional synthase-receptor pairs. The genes for synthase-receptor pairs are frequently co-located on the genome, and organisms can harbor more than one complete QS system (Fuqua et al., 1996). To increase the likelihood of identifying true pairs, we required that the two genes be separated by no more than two coding sequences. A total of 6,360 non-identical pairs were identified. We further discarded pairs that were more than 90% identical to another pair, resulting in 3,489 representative AHL synthase-receptor pairs.

We aligned these sequences to the LasI/R QS system from *P. aeruginosa* PAO1. Not only is *P. aeruginosa* a clinically important pathogen, the Las system is well-studied and crystal structures have been solved for both LasI (Gould et al., 2004) and LasR (Zou and Nair, 2009), making this a particularly useful model system for our studies. We connected the sequences of synthase and receptor from each pair and used GREMLIN to analyze covariation patterns in these sequences (Supplementary Fig. 1). We applied Average Product Correction (APC) to the GREMLIN covariance coefficients, a common technique shown to improve the accuracy of coevolution analyses (Buslje et al., 2009). We performed the same analysis by aligning the synthase-receptor pairs to the LuxI/R system from *Vibrio fischeri* MJ11. The top-ranking coevolving residue pairs overlap significantly between the LasI/R and LuxI/R systems (62.5% in common among the top 0.05% residue pairs) (Supplementary Fig. 1). We integrated the analyses for the LasI/R and LuxI/R systems by using the higher score for each residue pair and the top 10 residue pairs are shown in Fig. 2a. As a control, we randomly paired the synthases and receptors from different species and reanalyzed them using GREMLIN. While top-scoring covarying residues had a minimal GREMLIN score (with APC) of 0.09, the highest score from the randomized control was 0.08 (Fig. 2b). This control provides a guideline for our analysis; residues with a GREMLIN score (with APC) above or near the maximal score for the randomized control are likely to meaningful.

**Fig. 2.**
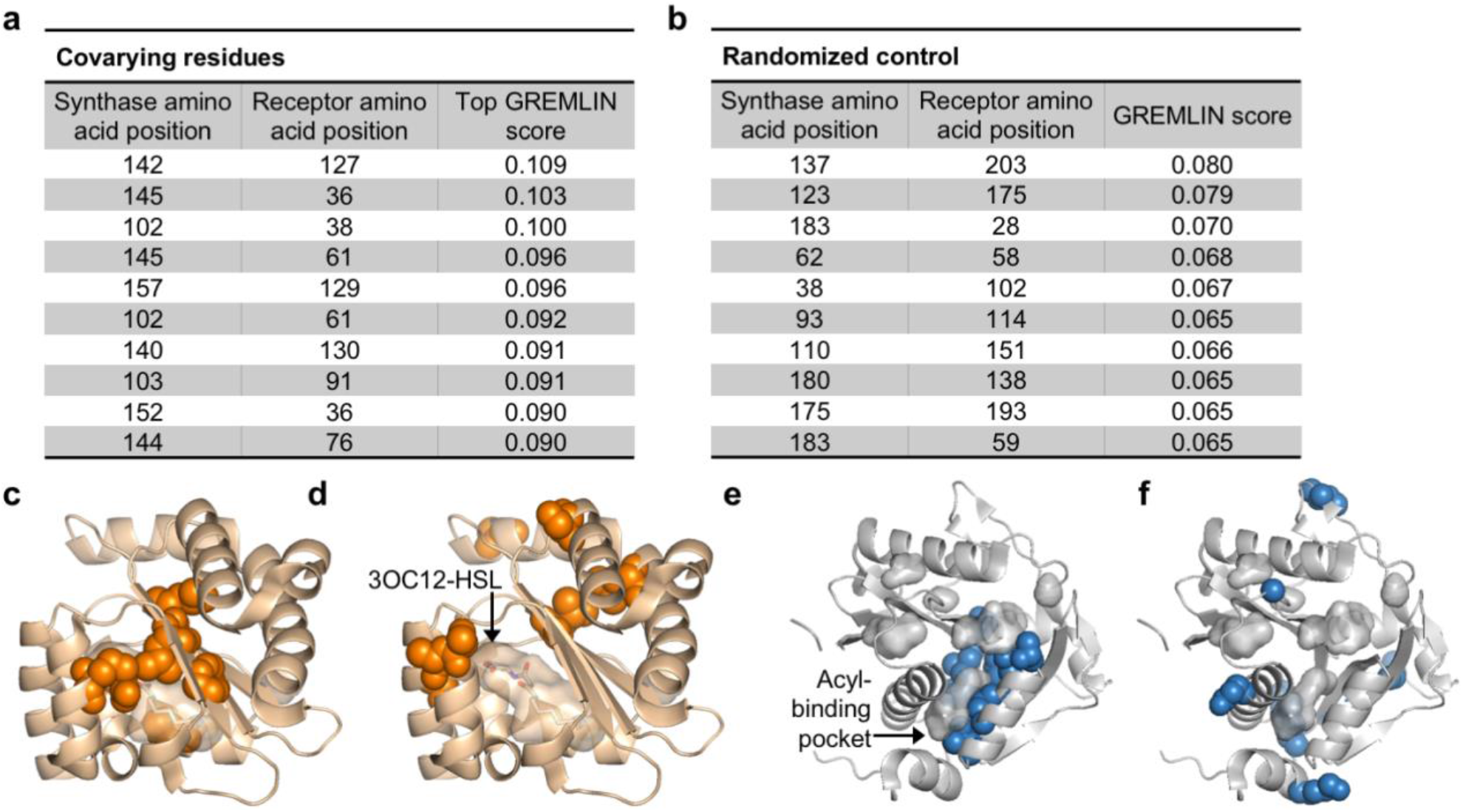
Covarying residues identified in LasI/R. **a)** Top-scoring covarying residues in LasI (synthase) and LasR (receptor) along with the top GREMLIN score (with APC) for each residue pair based an integrated analysis of the Las and Lux systems. **b)** Top-scoring residues in a randomized control, mapped onto LasI/R, along with the GREMLIN score (with APC) for each pair. **c)** Top-scoring covarying residues mapped onto LasR (covarying residues in orange, 3OC12-HSL in silver; PDB 3IX3) and **e)** LasI (covarying residues in blue; PDB 1RO5). Top-scoring residues in the randomized control are mapped onto **d)** LasR and **f)** LasI as in panels **c** and **e**.

### Top-scoring residues cluster near ligand-binding pockets

For both LasI and LasR, the top-scoring covarying residues cluster around the ligand-binding pocket. For LasR, the top-scoring residues map exclusively to the ligand-binding domain with an average distance of 5.0 Å from the co-crystalized native ligand *N*-3-oxo-dodecanoyl-L-homoserine lactone (3OC12-HSL) (Fig. 2c). In contrast, the residues identified in the randomized control are scattered throughout LasR, including three residues in the DNA-binding domain, and are an average distance of 17.8 Å from 3OC12-HSL (Fig. 2d).

In LasI, the top-scoring covarying residues cluster around the hydrophobic pocket thought to bind the fatty acyl substrate (Fig. 2e) and are an average distance of 3.7 Å from an acyl substrate modeled into the LasI structure (Gould et al., 2004). As with LasR, the residues identified in the randomized control are scattered throughout LasI, with many of the residues exposed to solvent (Fig. 2f). The randomized control residues in LasI are over three times further from the fatty acyl substrate, mean distance = 11.7Å, compared to the covarying residues.

Due to their location near the ligand-binding pockets, several of the covarying residues have been previously studied in various LasI/R homologues. Encouragingly, many of these residues have been reported to be important for protein activity and, in some cases, for selectivity. We have summarized several of these studies in Supplementary Tables 2 and 3.

### LasR mutations alter selectivity

To determine whether residues identified by GREMLIN are involved in LasR selectivity, we mutated a selection of the top-scoring amino acids, G38, R61, A127, S129, and L130, to the most common natural variants at each position (Supplementary Table 4). By expressing LasR in *Escherichia coli* and measuring its activity against a previously reported panel of 19 AHL signals (Wellington and Greenberg, 2019), we were able to quickly prioritize mutants for further study. The majority of our LasR mutants retained the ability to respond to AHLs and all mutants had an altered selectivity profile when compared to wild-type (Supplementary Fig. 2).

We, and others, have previously demonstrated that compared to native activity, QS receptor sensitivity and selectivity can be altered when in *E. coli* (Moore et al., 2015; Wellington and Greenberg, 2019). We therefore engineered several mutations into *lasR* on the *P. aeruginosa* PAO-SC4 chromosome to confirm our findings. *P. aeruginosa* PAO-SC4 is an AHL synthase-null mutant which we use here to measure LasR activity in response to exogenously provided AHL signals. The *lasR* mutations largely had the same effect on activity and selectivity in *P. aeruginosa* as they did when *lasR* was expressed *E. coli* (Supplementary Fig. 3). Of note, LasR^A127L^ had an increased sensitivity to numerous signals (Fig. 3a-d), potentially through increased hydrophobic interactions with the fatty acyl chain of the AHLs. Consistent with its role in a water-mediated hydrogen bond with the C3 oxygen of 3OC12-HSL, and with previous studies (Collins et al., 2006; Gerdt et al., 2015), LasR R61 mutants were less responsive to oxo-substituted AHLs, but maintained wild-type or better levels of activation by unsubstituted AHLs (Fig. 3a-d and Supplementary Fig. 3).

**Fig. 3.**
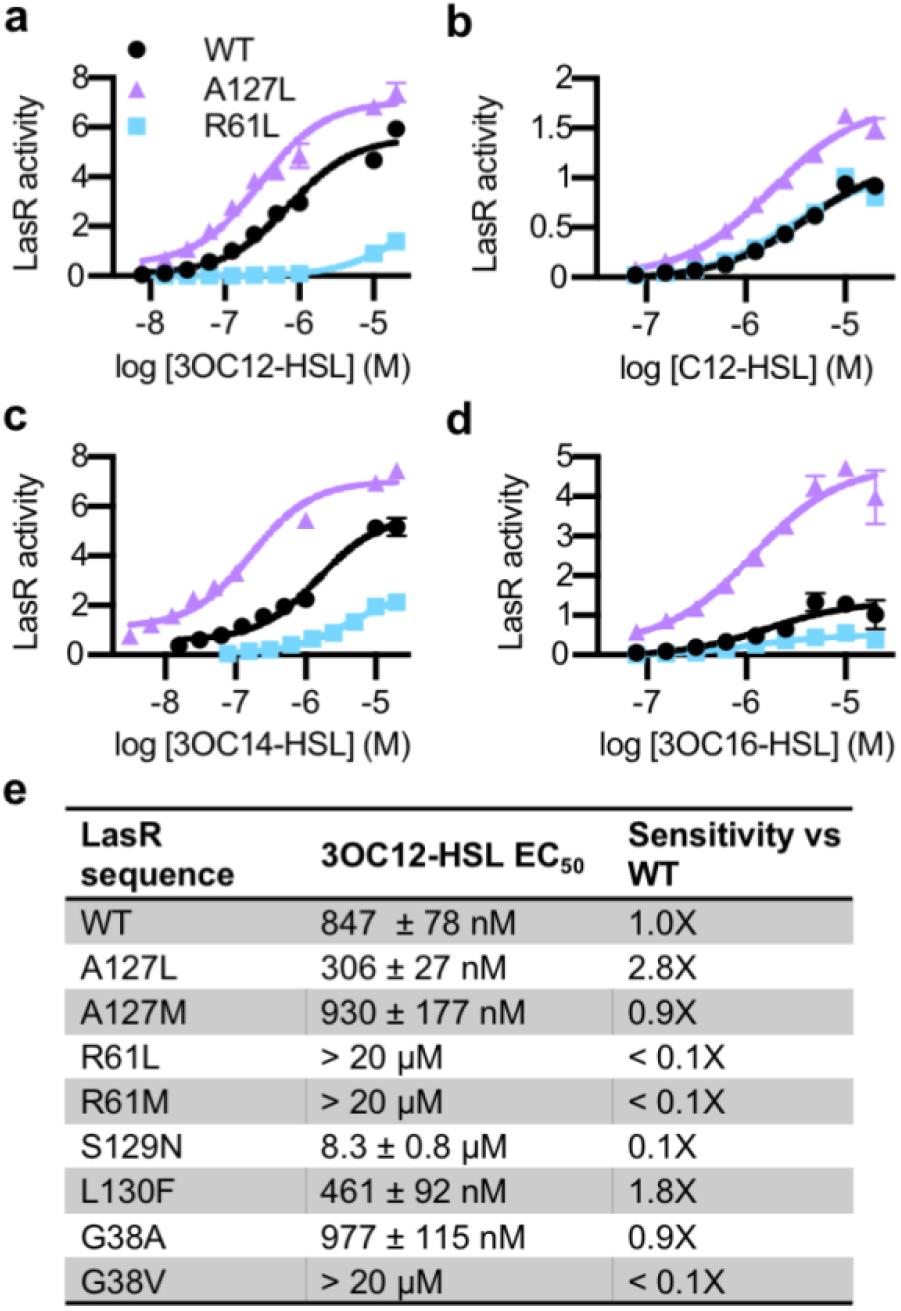
Activity of LasR mutants. Activity of chromosomal *lasR* mutants in *P. aeruginosa* PAO-SC4 pPROBE-P_rsaL_ in response to **a)** 3OC12-HSL, **b)** *N*-dodecanoyl-L-homoserine lactone (C12-HSL), **c)** *N*-3-oxo-tetradecanoyl-L-homoserine lactone (3OC14-HSL) or **d)** *N*-3-oxo-hexadecanoyl-L-homoserine lactone (3OC16-HSL). Indicated mutations are amino acid substitutions. Wild-type (WT) is shown in black, LasR^A127L^ in purple, and LasR^R61L^ in blue. The horizontal axis indicates AHL concentration. LasR activity is reported on the vertical axis as relative fluorescence units normalized by optical density at 600 nm (RFU/OD x 1,000). Data are the mean and standard deviation of three biological replicates and are representative of three independent experiments. **e)** Concentration for half-maximal activation (EC_50_) of 3OC12-HSL for *P. aeruginosa* PAO-SC4 LasR mutants calculated from data in Supplementary Fig. 3. Data are the mean and SEM of three (mutants) or four (wild-type) independent experiments. Sensitivity of the mutants compared to LasR^WT^ is calculated by dividing WT EC_50_ by mutant EC_50_.

The mutations also affected the sensitivity of LasR to 3OC12-HSL (Supplementary Fig. 3). Interestingly, two of our mutants were more sensitive to 3OC12-HSL than wild-type LasR. LasR^A127L^ was roughly 3-fold more sensitive and LasR^L130F^ was 2-fold more sensitive (Fig. 3e). This increased sensitivity came at the cost of decreased selectivity for both of these mutations. In fact, many of our single amino acid mutants displayed reduced selectivity compared to wild-type LasR (Supplementary Fig. 2 and Supplementary Fig. 3).

### LasI mutations alter activity and selectivity

Similar to LasR, we focused our LasI mutations on the top-scoring positions: L102, T142, T145, and L157 (Supplementary Table 5). We expressed wild-type or mutated *lasI* on a low copy number plasmid in the AHL synthase-null *P. aeruginosa* PAO-SC4 and extracted AHLs produced by these bacteria from culture fluid. While bioassays are commonly used for the detection of AHLs (Chu et al., 2011), they suffer from multiple drawbacks. In particular, bioassays are not equally sensitive to all AHLs and typically cannot be used to determine which AHLs are produced and in what ratio. To screen our LasI mutants for altered activity and selectivity, we developed a thin layer chromatography (TLC) method based on our existing high performance liquid chromatography (HPLC) radiotracer assay (Schaefer et al., 2018). In this method, the C1 position in the homoserine lactone ring is labeled with ^14^C. The label is incorporated into AHLs at a ratio of one ^14^C per AHL molecule. This results in unbiased detection of all AHLs produced. While the established method uses HPLC to separate and detect AHLs one sample at a time, we can run nine samples per TLC, resulting in a more high-throughput assay.

Using our TLC method, we confirmed that *lasI* directs the synthesis of the same primary product whether it is expressed on a plasmid or from the chromosome (Supplementary Fig. 4). HPLC analysis of matched extracts confirmed that the major LasI product observed by TLC is 3OC12-HSL. As expected, an empty vector control did not produce detectable AHLs, nor did we detect radioactivity in a media-only control. We then screened the activity each *lasI* mutant by TLC (Supplementary Fig. 4). Several mutants produced little or no detectable AHLs, while some appeared to produce more 3OC12-HSL than wild-type LasI. We analyzed select extracts by both TLC and HPLC and found that the results were consistent between the two methods, further validating the TLC method.

Based on our TLC results we selected one mutant, LasI^L157W^, for further study by HPLC. We found that LasI^L157W^ produces equal amounts of two ^14^C-AHLs that elute in the fractions of *N*-3-oxo-decanoyl-L-homoserine lactone (3OC10-HSL) and *N*-3-oxo-octanoyl-L-homoserine lactone (3OC8-HSL) along with a lesser amount of 3OC12-HSL (Fig. 4). These findings demonstrate that the covarying residues influence LasI activity and selectivity, and that a single mutation is sufficient to significantly alter LasI selectivity.

**Fig. 4.**
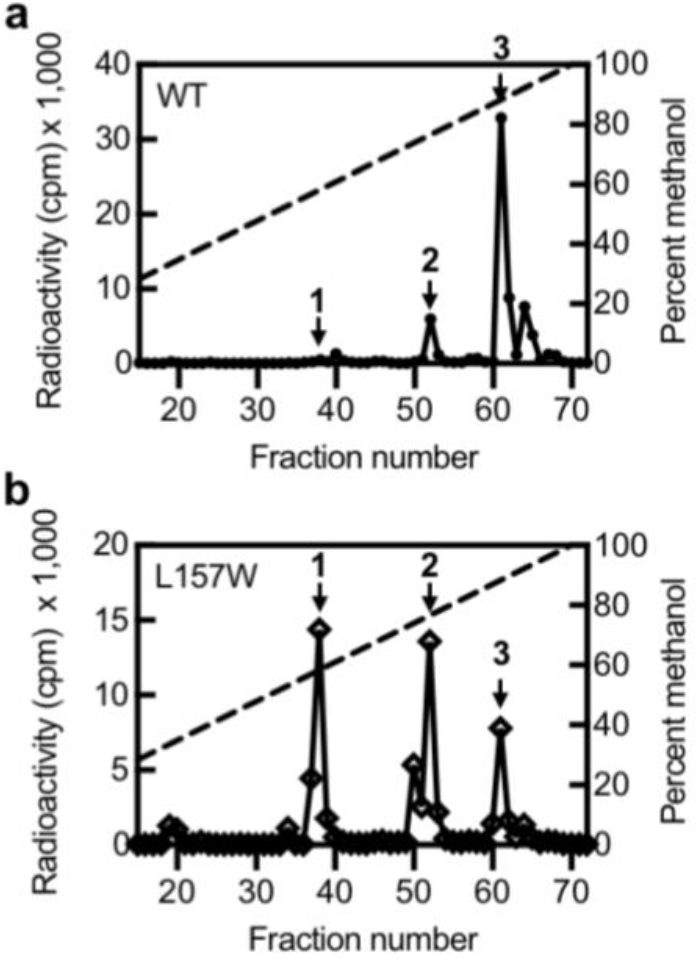
Activity of LasI mutants. HPLC analysis of radiolabeled AHLs extracted from *P. aeruginosa* PAO-SC4 harboring **a)** pJN-RBSlasI^WT^ or **b)** pJN-RBSlasI^L157W^. The horizontal axis denotes the HPLC fraction number (fractions 1-14 are not shown). The methanol gradient is indicated as a dashed line plotted on the right vertical axis. The left vertical axis indicates the amount of radioactivity (counts per minute (cpm)) in each fraction. Data are representative of two (L157W) or three (WT) independent experiments. Arrow 1 indicates the fraction in which 3OC8-HSL elutes, arrow 2 indicates the fraction in which 3OC10-HSL elutes, and arrow 3 indicates the fraction in which 3OC12-HSL elutes.

### Multiple mutations can “rewire” LasI/R selectivity

In general, multiple mutations are required to generate a protein with orthogonal selectivity (Aakre et al., 2015; Collins et al., 2006; Skerker et al., 2008). In non-QS proteins, altered selectivity has been engineered by swapping the covarying residues in one homolog to the identities in another (Aakre et al., 2015; Skerker et al., 2008). Here, we seek to “rewire” LasI/R to use an orthogonal signal. We targeted the MupI/R system from *Pseudomonas fluorescens* NCIMB 10586, which uses the signal 3OC10-HSL (Hothersall et al., 2011). MupI and MupR share 52% and 39% identity with LasI and LasR, respectively.

LasR and MupR differ at eight covariation sites in the ligand-binding domain with a GREMLIN score (with APC) > 0.08 (Supplementary Fig. 5). LasR modified to contain all eight mutations was inactive. However, there were several intermediate mutants that displayed an increased response to 3OC10-HSL. We identified three mutations that are sufficient for this increased sensitivity: L125F, A127M, and L130F (Fig. 5a). LasR^L125F, A127M, L130F^ is over 20-fold more sensitive to 3OC10-HSL than wild-type LasR. The L125F mutation appears to be the primary driver of this altered selectivity (Fig. 5b,c and Supplementary Fig. 5). All “MupR-like” LasR mutants responded to 3OC12-HSL with similar sensitivity to wild-type LasR (Supplementary Fig. 5).

**Fig. 5.**
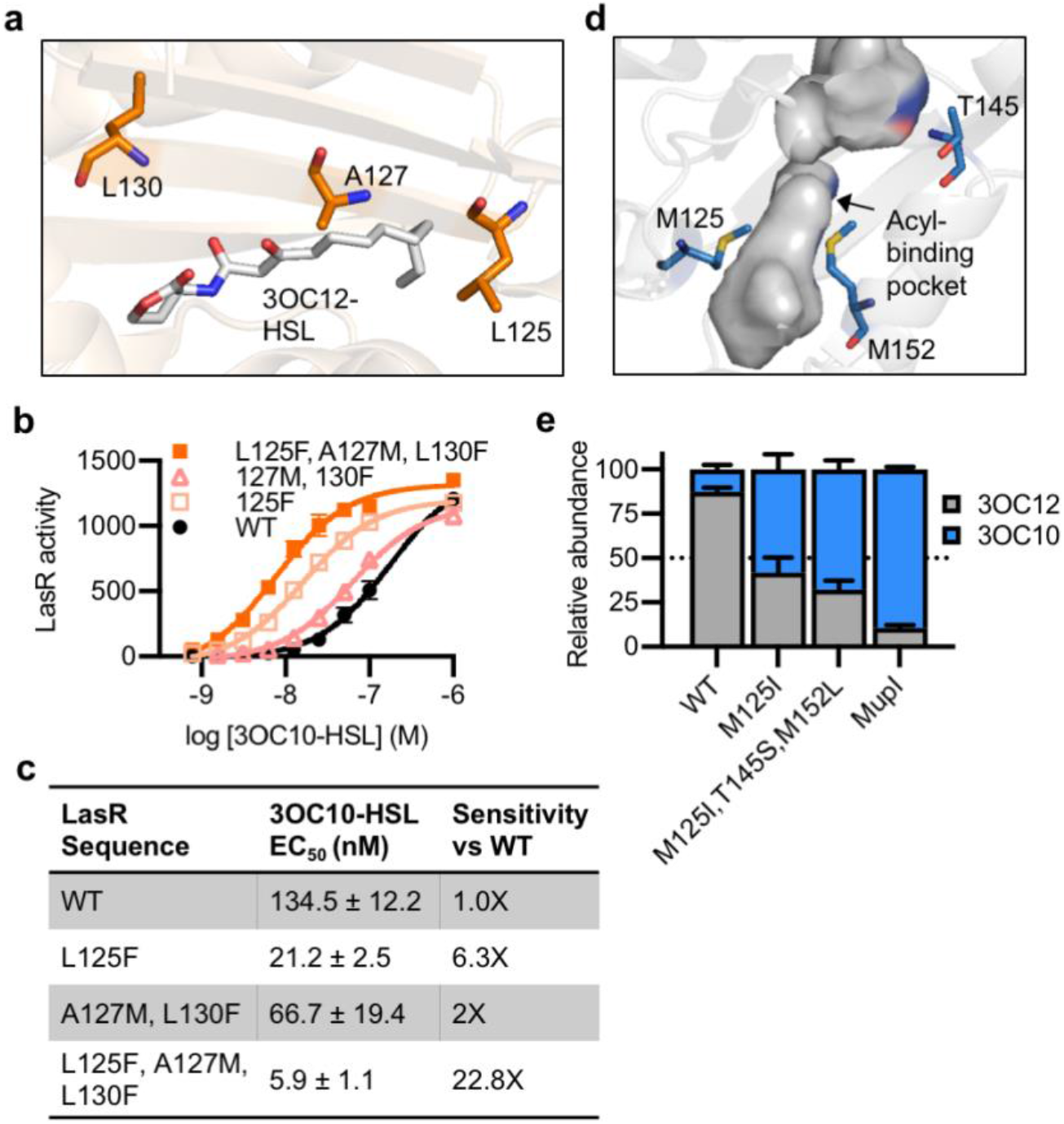
Reprogramming LasI/R selectivity. **a)** Residues mutated in LasR shown as orange sticks in the LasR structure (PDB 3IX3). 3OC12-HSL is shown in grey. **b)** LasR activity in response to 3OC10-HSL measured in *E. coli* harboring pJNL (wild-type, WT, or with indicated mutations) and pPROBE-PrsaL. Data are the mean and standard deviation of three biological replicates and are representative of three independent experiments. **c)** Concentration of half maximal activity (EC_50_) of 3OC10-HSL for LasR, calculated from data shown in panel **b**. Data are mean and SEM. Sensitivity of mutants compared to LasR^WT^ is calculated by dividing the WT EC_50_ by mutant EC_50_. **d)** Residues mutated in LasI shown as blue sticks in the LasI structure (PDB 1ROH). **e)** Relative amount of AHLs produced by *P. aeruginosa* PAO-SC4 harboring pJN-RBSlasI (WT or with the indicated mutations) or pJN-RBSmupI. Ratios were calculated from HPLC data shown in Supplementary Fig. 6. Bars show mean and standard deviation. The dashed line indicates equal production of 3OC10-HSL and 3OC12-HSL.

LasI differs from MupI at five high-scoring covariation residues: LasI M125, T145, M152, V159, and N181 (Supplementary Fig. 6), the first 3 of which line the LasI acyl-binding pocket (Fig. 5d). Swapping these three residues for their MupI identities resulted in a synthase that has substantially altered selectivity. LasI^M125I, T145S, M152L^ produces ~2-fold more 3OC10-HSL than 3OC12-HSL (Fig. 5e). The M125I mutation alone was sufficient to relax LasI’s selectivity, resulting in a synthase that produces roughly equal amounts of 3OC10-HSL and 3OC12-HSL. As a comparison, we measured the activity of *mupI* expressed in *P. aeruginosa*, and found it produces 9:1 3OC10-HSL:3OC12-HSL (Fig. 5e and Supplementary Fig. 6). All single and double “MupI-like” LasI mutants retained AHL synthase activity, but only those mutants that contain the M125I mutation displayed increased 3OC10-HSL production relative to 3OC12-HSL (Supplementary Fig. 6).

## Discussion

Despite decades of study, it has been challenging to determine how AHL QS systems distinguish between signals. We hypothesized that we could identify covariation patterns in AHL QS systems and that these patterns would illuminate residues important for signal selectivity. By analyzing the sequences of 6,360 unique QS systems, we identified amino acids that strongly covary between AHL synthases and receptors. The top-scoring residues in our analysis cluster near the ligand-binding pockets for both proteins and are more than three times closer to the signal molecule compared to top-scoring residues in a randomized control. We focused our study on *P. aeruginosa* LasI/R. Through targeted mutations in the top-scoring covarying residues we demonstrate that these amino acids are indeed determinants of signal selectivity. We have thus validated a new application of covariation analysis for proteins that interact indirectly and not through direct binding to one-another. Additionally, these strong covariation results further support the view that AHL synthases and receptors coevolve.

For both the synthase, LasI, and the receptor, LasR, a single amino acid substitution is sufficient to significantly alter selectivity. Interestingly, our mutations also revealed that LasR is not optimized to be as sensitive as possible to its native 3OC12-HSL signal. The increase in sensitivity of specific mutants came at the cost of decreased selectivity, which suggests that QS systems may evolve to balance these two properties. Furthermore, increased sensitivity to the native signal may lead to premature activation of the QS regulon, which would likely decrease fitness (Darch et al., 2012). The mutants generated in our study provide us with the tools to directly address these questions and assess the impact of sensitivity and selectivity on QS function.

We also demonstrated that we can use covarying residues to rationally engineer a QS system to produce and respond to a signal of our choosing. By mutating the covarying residues in LasI/R, we improved the sensitivity of LasR to 3OC10-HSL over 20-fold and increased the production of 3OC10-HSL by LasI roughly 15-fold. For both the synthase and receptor, a single amino acid substitution was the primary driver of the altered selectivity. This was surprising given the low sequence identity between LasI/R and MupI/R. These findings suggest new QS systems might evolve with relative ease. Further, the ability to engineer QS selectivity could be beneficial to synthetic biology where AHL signaling is a powerful tool to build biological circuits (Davis et al., 2015).

Though we were able to substantially increase the 3OC10-HSL activity of LasI/R, our mutants retained their native 3OC12-HSL activity. We have thus generated a promiscuous system with broadened selectivity. Similarly, a directed evolution study of the AHL receptor LuxR found that it evolves through promiscuous intermediates (Collins et al., 2005). This has also been observed in other systems, such as toxin-antitoxin systems (Aakre et al., 2015). Proteins tend to evolve through broadly active intermediates before gaining new specificity. In this way, the system maintains functionality *en route* to altered selectivity. Quorum sensing systems appear to follow these same trends.

One limitation we faced is a lack of close LasI/R homologs with known signals. It has been demonstrated that “supporting” residues, i.e. residues within a protein that covary with the selectivity residues, may indirectly impact selectivity by influencing the orientation of selectivity residues (Aakre et al., 2015). Thus, given the large differences in sequence identity between the Las and Mup systems, there are likely other residues that must be mutated to fully swap selectivity. The identification of a more closely related system to LasI/R may provide a better starting point for engineering altered selectivity. Alternatively, our mutants could be further evolved though saturating mutagenesis and/or *in vitro* evolution.

Collectively, our results provide insight into AHL QS selectivity and will help us predict signal selectivity in newly identified QS systems, in metagenomes, and in naturally occurring QS variants such as those found in clinical isolates. More broadly, we have gained insight into how AHL QS systems evolve and diversify.

## Methods

### Identification of quorum sensing systems

Starting from 24 pairs of manually curated QS synthases and receptors (Supplementary Table 1), we searched for homologs in complete bacterial genomes using BLAST (e-value <0.01) (Altschul et al., 1990). We filtered the BLAST hits by sequence identity (>30%) to the query and the alignment coverage (query coverage >0.75 and hit coverage >0.75), and the filtered hits were aligned by Clustal Omega (Sievers and Higgins, 2021). We selected the LasI/R system from *Pseudomonas aeruginosa* PAO1 as the target and mapped the Multiple Sequence Alignments (MSA) to the target system. We built sequence profiles from the MSA with HMMER (Eddy, 2009) and hmmbuild for the QS synthases and receptors, respectively. The sequence profiles were then used to search against the European Nucleotide Archive database (Amid et al., 2019) and the Integrated Microbial Genomes and Microbiomes database (Chen et al., 2020) from Joint Genome Institute using HMMER hmmsearch. A total of 149,837 and 5,046,620 homologs were found in these databases for the QS synthase and receptor, receptively. Because the synthases and receptors of the known QS systems frequently locate near each other in the genome, we kept synthase-receptor gene pairs that are separated by no more than two other open reading frames (ORFs) in the genome or contig. A total of 14,980 synthase-receptor gene pairs were identified and they represent 6,360 non-identical QS systems. In another attempt, we carried out the same procedure using the LuxI/R system from *Vibrio fischeri* MJ11 as the target system. A similar number of QS systems were identified.

### Identification of covarying residues

We connected the synthase and receptor protein sequences for each QS system we found in the databases and derived the alignments between these QS systems to the target QS system (LasI/R) from the hmmsearch result. We filtered the MSA for synthase-receptor pairs by sequence identity (maximal identify for remaining sequences <=90%) and gap ratio in each sequence (maximal gap ratio <=25%). We applied GREMLIN to analyze the covariation in the MSA (Kamisetty et al., 2013), and the GREMLIN coefficients were normalized using Average Product Correction (APC) (Buslje et al., 2009) as we described previously(Ovchinnikov et al., 2014). The GREMLIN coefficients after APC were used as measures for covariation signals between synthase and receptor amino acid residues. As a control, we connected each synthase sequence with a randomly selected receptor sequence and performed the covariation analysis in the same way.

We mapped the top-scoring covarying residues in the LasI/R system onto the crystal structures for each protein. Reported distances between residues and ligands are the shortest distance between any non-hydrogen atoms. For LasR, distances were calculated using PDB 6V7X. For LasI, distances were calculated using a LasI structure with 3-oxo-C12-acyl-phosphopantetheine modeled into the acyl-binding pocket (Gould et al., 2004). Reported distances for LasI are between residues and the acyl portion of the modeled substrate.

### Bacterial strains, plasmids, and culture conditions

Bacterial strains and plasmids are listed in Supplementary Table 6. Unless otherwise specified, *Pseudomonas aeruginosa* and *Escherichia coli* were grown in lysogeny broth (LB) (10 g tryptone, 5 g yeast extract, 5 g NaCl per liter) buffered with 50 mM 3-(*N*-morpholino) propanesulfonic acid (MOPS) (pH 7) (LB/MOPS) or on LB agar (LB plus 1.5% Bacto agar) (Wellington and Greenberg, 2019). Liquid cultures were grown at 37°C with shaking. For radiotracer thin layer chromatography (TLC) experiments, *P. aeruginosa* was grown in Jensen’s medium with 0.3% glycerol (Schaefer et al., 2018).

For plasmid selection and maintenance, antibiotics were used at the following concentrations: *P. aeruginosa*, 30 μg per mL gentamicin (Gm) and 150 μg per mL carbenicillin (Cb); *E. coli* 10 μg per mL Gm and 100 μg per mL ampicillin (Ap). BD Difco Pseudomonas Isolation Agar (PIA) was prepared according to manufacturer directions and supplemented with 100 μg per mL Gm as needed. Where needed for gene expression L-arabinose (0.4% w/v) was added.

All chemicals and reagents were obtained from commercial sources. AHLs were dissolved either in dimethyl sulfoxide (DMSO) or in ethyl acetate (EtAc) acidified with glacial acetic acid (0.01% v/v). AHLs in DMSO were used at <=1% of the final culture volume and AHLs dissolved in EtAc were dried on the bottom of the culture vessel prior to addition of the bacterial culture. DMSO or acidified EtAc was used as a vehicle control where appropriate.

### Plasmid and strain construction

pJN-lasI and pJN-RBSmupI were constructed using *E. coli*-mediated DNA assembly (Kostylev et al., 2015). Briefly, for pJN-lasI, *lasI* was amplified from *P. aeruginosa* PAO1 genomic DNA (gDNA) using primers lasI-pJN-F and lasI-pJN-R (Supplementary Table 6). pJN105 was amplified using the reverse complement of these primers. The resulting PCR products were treated with the restriction enzyme DpnI to remove the parent template. Both PCR products were then used to transform *E. coli* (NEB 5alpha). The resulting constructs were confirmed by Sanger sequencing. For pJN-RBSmupI, we began by amplifying *mupI* from *Pseudomonas fluorescens* Migula (ATCC 49323) gDNA using primers mupI-F and mupI-R. We then used primers mupI-pJN-F and mupI-pJN-R to amplify the *mupI* PCR product and used the reverse complement of these two primers to amplify pJN-RBSlasI. The resulting PCR products were treated the same as for pJN-lasI. We constructed pJN-RBSlasI using restriction digestion. The *lasI* gene, including its upstream ribosomal binding site (RBS), was amplified from *P. aeruginosa* PAO1 gDNA using primers RBS-lasI-F and lasI-pJN-R. pJN-lasI and the RBS-*lasI* PCR product were digested using NheI and SacI, gel or column purified respectively, ligated by T4 DNA ligase, and transformed into NEB 5alpha. The resulting constructs were confirmed by Sanger sequencing. Plasmids were introduced into *E. coli* by using heat shock and were introduced into *P. aeruginosa* by electroporation.

Point-mutations were introduced to *lasI* and *lasR* on pJN-lasI and JNL or pEXG2-lasR, respectively, using site directed mutagenesis by PCR. Primers were designed to amplify each plasmid while introducing the desired mutation(s). The resulting PCR products were treated with DpnI and were then used to transform NEB 5alpha. Plasmids from the resulting colonies were screened for the desired mutations by Sanger sequencing. To mutate *lasR* on the *P. aeruginosa* PAO-SC4 chromosome, *E. coli* S17-1 was used to deliver pEXG2-lasR containing various *lasR* mutations to PAO-SC4 via conjugation and potential mutants were isolated as previously described(Kostylev et al., 2019). All mutations were confirmed by PCR amplification of *lasR* from the genome followed by Sanger sequencing.

### LasR activity measurements

LasR activity was measured in *E. coli* containing pJNL and pPROBE-P_rsaL_ or in *P. aeruginosa* PAO-SC4 containing pPROBE-P_rsaL_ using previously reported methods (Wellington and Greenberg, 2019). Briefly, overnight-grown cultures were diluted 1:100 and grown back to log-phase. For *E. coli*, cultures were grown to an optical density at 600 nm (OD) of 0.3, treated with L-arabinose (0.4%), and incubated with AHLs for 4 h. For *P. aeruginosa*, cultures were grown to an OD between 0.05 and 0.3, were diluted to an OD of 0.01 and then incubated with AHLs for 16 to 18 h. LasR activity was measured as GFP fluorescence (excitation 490 nm, emission 520 nm, gain 50) using a Synergy H1 microplate reader (Biotek Instruments). Activity measurements were normalized by dividing by OD and subtracting background values (fluorescence per OD for cultures incubated with vehicle control). Concentrations of half maximal activation, EC_50_, were calculated using GraphPad Prism.

### TLC screening for AHLs

Cultures of *P. aeruginosa* PAO1Δ*rhlI* or of *P. aeruginosa* PAO-SC4 with pJN-empty or with wild-type or mutated pJN-lasI were grown overnight in Jensen’s medium with 0.3% glycerol. Overnight cultures were used to inoculate fresh medium (1% v/v). When the OD reached 0.5, *lasI* expression was induced with arabinose (0.4%) and 1.1 mL cultures were incubated with 1.1 μCi/mL L-[1-^14^C]-methionine (^14^C-methionine) for 90 min (Schaefer et al., 2018). Cells were pelleted by centrifugation and 1 mL of supernatant fluid was extracted twice with 2 mL acidified EtAc. The extracts were dried under N_2_ and resuspended in 15 μL acidified EtAc. Five μL of each extract was spotted on an aluminum backed C18-W-silica TLC plate (Sorbtech). AHLs were separated using 70% methanol in water, then the TLC plate was dried and exposed to a phosphor screen for at least 16 h. Phosphor screens were imaged with a Sapphire Biomolecular Imager (Azure Biosystems). To confirm TLC findings, select extracts were dried, suspended in methanol and analyzed by C18-reversed-phase high performance liquid chromatography (HPLC) using a previously reported method (Schaefer et al., 2018).

### HPLC radiotracer assays for LasI activity

For better detection of AHLs, we slightly modified the radiolabeling protocol detailed above, modeling it after a previously published method (Leadbetter and Greenberg, 2000). Cultures of *P. aeruginosa* PAO-SC4 with wild-type or mutated pJN-RBSlasI were grown overnight in LB/MOPS. Overnight cultures were used to inoculate 5 mL LB/MOPS (1% v/v). After 2 h, *lasI* expression was induced with arabinose (0.4%) and cultures were grown to OD 0.7. Cells were centrifuged at 5,000 rpm for 10 min, and pellets were suspended in 1.1 mL phosphate buffered saline (PBS) with 10 mM glucose. After shaking incubation at 37°C for 10 min, 1.1 μCi ^14^C-methionine was added to the cell suspension. Cell suspensions were incubated with radiolabel for 2 h, after which cells were pelleted by centrifugation and 1 mL supernatant fluid was extracted twice with 2 mL acidified EtAc. Radiolabeled AHLs were dried under N_2_ and suspended in methanol. One-third of each extract was analyzed by reversed-phase HPLC using a gradient of 10 to 100% methanol-in-water (Schaefer et al., 2018).

## Acknowledgements

We thank Mair Churchill for sharing her lab’s LasI structure modeled with an acyl substrate. This work was supported by grants NIH R35GM136218 to EPG, Yeast Program Grant 5 P41 GM103533-24, Washington Research Foundation fellowship to QC, and Helen Hay Whitney Foundation fellowship to SWM.

## Author contributions

SWM, QC, DB, and EPG conceived of and designed the investigation. QC carried out bioinformatic analyses. SWM, EM, and AZ constructed mutants and measured their activity. SWM and ALS developed and conducted radioassays. SWM together with QC, ALS, EPG, and DB interpreted data and wrote the manuscript.

