## Supplementary material for "A covariation analysis reveals elements of selectivity in quorum sensing systems"

**Supplementary Table 1 | Manually curated QS synthases and receptors.**

| QS system name | Bacterium | Cognate signal | GenBank (I/R) |
|---|---|---|---|
| LasI/R[a] | *Pseudomonas aeruginosa* PAO1 | 3OC12-HSL | AAG04821.1/ AAG04819.1 |
| RhlI/R | *P. aeruginosa* PAO1 | C4-HSL | AAG06864.1/ AAG06865.1 |
| PfvI/R | *Pseudomonas fuscovaginae* UPB0736 | 3OC10/ 3OC12-HSL | CAQ15950.2/ CAQ15948.2 |
| PfsI/R | *P. fuscovaginae* UPB0736 | C10/ C12-HSL | CBI67625.1/ CBI67623.1 |
| CepI/R | *Burkholderia vietnamiensis* G4 | C8-HSL | ABO58211.1/ ABO58209.1 |
| BviI/R | *B. vietnamiensis* G4 | C10-HSL | ABK32009.1/ ABK32010.1 |
| BtaI/R1 | *Burkholderia thailandensis* E264 | C8-HSL | ABC35524.1/ ABC34804.1 |
| BtaI/R2 | *B. thailandensis* E264 | 3OHC10-HSL | ABC34067.1/ ABC34774.1 |
| BtaI/R3 | *B. thailandensis* E264 | 3OHC8-HSL | AIP27810.1/ AIP27980.1 |
| CepI/R | *Burkholderia cenocepacia* J2315 | C8-HSL | CAR55728.1/ CAR55726.1 |
| AbaI/R | *Acinetobacter baumannii* ATCC17978 | 3OHC12-HSL | AKQ28471.1/ AKQ28469.1 |
| CviI/R | *Chromobacterium violaceum* ATCC 12472 | 3OHC10-HSL | AAQ61751.2/ AAQ61750.2 |
| CviI/R | *Chromobacterium violaceum* ATCC 31532 | C6-HSL | PLV42917.1/ ADC79709.1 |
| LuxI/R | *Vibrio fischeri* ES114 | 3OC6-HSL | AAW87994.1/ AAW87995.1 |
| LuxI/R[a] | *Vibrio fischeri* MJ11 | 3OC6-HSL | ACH64323.1/ ACH63788.1 |
| ExpI/R | *Pectobacterium parmentieri* SCC3193 | 3OC8-HSL | AFI92653.1/ AFI92652.1 |
| BjaI/R | *Bradyrhizobium japonicum* USDA110 | Isovaleryl-HSL | BAC46328.1/ BAC46327.1 |
| MupI/R | *Pseudomonas fluorescens* NCIMB 10586 | 3OC10-HSL | AAK28505.1/ AAK28504.1 |
| PpuI/R | *Pseudomonas putida* IsoF | 3OC10/3OC12-HSL | AAM75411.1/ AAM75413.1 |
| TraI/R | *Agrobacterium tumefaciens* C-58 | 3OC8-HSL | AAK91000.1/ AAK91098.1 |
| MbaI/R | *Methylobacter tundripaludum* 21/22 | 3OHC10-HSL | WP_150113271.1/ WP_006890625.1[b] |
| YruI/R | *Yersinia ruckeri* ATCC 29473 | 3OC8-HSL | KGA49182.1/ KGA49159.1 |
| GtaI/R | *Rhodobacter capsulatus* SB1003 | C16-HSL | ADE84094.1/ ADE84093.1 |
| AhyI/R | *Aeromonas hydrophila* ML09-119 | C4-HSL | AGM42354.1/ AGM42355.1 |

[a] Covariation analyses were mapped onto LasI/R from *P. aeruginosa* PAO1 and LuxI/R from *V. fischeri* MJ11.

[b] The accession number provided is for the NCBI Reference Sequence.

**Supplementary Table 2 | Select previously reported data for LasR homologs with relevant mutations**. The native signal for each receptor is indicated in parentheses.

| LasR residue | LasR homolog | Mutation | Effect on receptor activity[a] | Reference |
|---|---|---|---|---|
| **L36** | LuxR (3OC6-HSL) | L42A[b]<br>L42S | Reduced sensitivity<br>Reduced sensitivity | (Koch et al., 2005) |
| | TraR[c] (3OC8-HSL) | A38V | Reduced activity | (Luo et al., 2003) |
| **G38** | QscR (3OC12-HSL) | G40F | Altered selectivity: impaired response to 3OC12-HSL, increased sensitivity to 3OC6-HSL | (Lintz et al., 2011) |
| **P57** | TraR | H54Y | Reduced activity | (Luo et al., 2003) |
| **R61** | LasR (3OC12-HSL) | R61M | Altered selectivity: impaired response to 3OC12-HSL, maintained response to C12-HSL | (Collins et al., 2006) |
| | LasR | R61M | Reduced sensitivity | (Gerdt et al., 2015) |
| | LuxR | R67M | Altered selectivity: impaired response to 3OC6-HSL, maintained response to C6-HSL | (Collins et al., 2006) |
| | TraR[c] | Q58L | Altered selectivity: improved sensitivity to 3OC6-HSL | (Chai and Winans, 2004) |
| | | Q58F | Reduced activity | |
| **T75** | LasR | T75V | Increased sensitivity | (Gerdt et al., 2015) |
| **V76** | QscR | V78F | Reduced activity | (Lintz et al., 2011) |
| **A127** | LasR | A127W | Altered selectivity: impaired response to 3OC12-HSL, maintained response to shorter AHLs | (McCready et al., 2019b) |
| | LasR | A127F | Reduced sensitivity | (McCready et al., 2019a) |
| | LuxR | M135I<br>M135V | Altered selectivity: impaired response to 3OC6-HSL; improved response to C8-HSL | (Collins et al., 2005) |
| | LuxR | M135A | Reduced sensitivity | (Koch et al., 2005) |
| **S129** | LasR | S129A | Reduced sensitivity | (Gerdt et al., 2015; Manson et al., 2020) |
| | LasR | S129C<br>S129W<br>S129F<br>S129T<br>S129M | Reduced sensitivity<br>Reduced sensitivity<br>Reduced sensitivity<br>Reduced sensitivity<br>Reduced sensitivity | (McCready et al., 2019b) |
| | TraR | T129S | No change from wild-type | (Chai and Winans, 2004) |
| | | T129L | Reduced activity | |
| | | T129I | Reduced activity | |
| | | T129F | Reduced activity | |
| | | T129A<br>T129V | Altered selectivity: impaired activity; equally sensitive to C8-HSL and 3OC8-HSL | |
| **L130** | LasR | L130F | Altered selectivity: increased sensitivity to 3OC12-HSL and to other AHLs | (McCready et al., 2019b) |

[a] All studies measured the activity of the receptor in *Escherichia coli* unless otherwise noted.

[b] Numbering is according to the *Vibrio fischeri* ES114 sequence.

[c] This study was performed in *Agrobacterium tumefaciens*, using a *traR* expression plasmid.

**Supplementary Table 3 | Select previously reported data for LasI homologs with relevant mutations.** The native signal for each synthase is indicated in parentheses.

| LasI residue | LasI homolog | Mutation | Effect on synthase activity[a] | Reference |
|---|---|---|---|---|
| **S103** | RhlI (C4-HSL) | S103E | Impaired activity | (Parsek et al., 1997) |
| | EsaI (3OC6-HSL) | S99A | Impaired activity | (Watson et al., 2002) |
| **T142** | LasI (3OC12-HSL) | T142G<br>T142A<br>T142S | Impaired activity<br>Slightly altered selectivity<br>Slightly altered selectivity | (Gould et al., 2006) |
| | EsaI (3OC6-HSL) | T140A | Altered selectivity: Increased production of C6-HSL | (Gould et al., 2006; Watson et al., 2002) |
| | EsaI | T140V | Impaired activity | (Watson et al., 2002) |
| **T144** | LasI | T144V | Impaired activity | (Gould et al., 2006) |
| **M152** | RhlI | F147L[b] | Increased activity | (Kambam et al., 2009) |
| | MesI[c] (C6-HSL) | L153A | Altered selectivity: Increased C8-HSL production | (Dong et al., 2020) |
| | | L153F | Altered selectivity: Increased C4-HSL production | |
| | BjaI[c] (isovaleryl-HSL) | F147Y | Altered selectivity: Increased C4-HSL production | (Dong et al., 2020) |

[a] All studies measured activity of the synthase in *Escherichia coli* unless otherwise noted.

[b] F147L was one of three amino acid substitutions in a synthase obtained by directed evolution.

[c] Substrate selectivity was studied using purified enzymes.

**Supplementary Table 4 | Most common amino acids at selected positions in LasR homologs.** Residue numbering according to LasR.

| Residue in receptor | Amino acid | Relative abundance |
|---|---|---|
| 38 | Gly* | 0.482 |
| | Leu | 0.095 |
| | Val | 0.086 |
| | Ala | 0.079 |
| 61 | Val | 0.231 |
| | Gln | 0.160 |
| | Leu | 0.128 |
| | Met | 0.126 |
| | Arg* | 0.093 |
| 127 | Leu | 0.287 |
| | Met | 0.174 |
| | . . . | . . . |
| | Ala* | 0.102 |
| 129 | Ser* | 0.536 |
| | Thr | 0.273 |
| | Ala | 0.092 |
| | Asn | 0.049 |
| 130 | Leu* | 0.359 |
| | Phe | 0.253 |
| | Ile | 0.176 |

* Indicates identity in wild-type LasR.

**Supplementary Table 5 | Most common amino acids at selected positions in LasI homologs.** Residue numbering according to LasI.

| Residue in synthase | Amino acid | Relative abundance |
|---|---|---|
| 102 | Leu* | 0.398 |
| | Ser | 0.157 |
| | Met | 0.156 |
| | Ile | 0.094 |
| 142 | Gly | 0.379 |
| | Thr* | 0.314 |
| | Ala | 0.154 |
| 145 | Ser | 0.231 |
| | Thr* | 0.199 |
| | Pro | 0.170 |
| | Asp | 0.135 |
| | . . . | . . . |
| | Ala | 0.009 |
| 157 | Trp | 0.271 |
| | Val | 0.220 |
| | Phe | 0.171 |
| | Leu* | 0.124 |

* Indicates identity in wild-type LasI.

**Supplementary Table 6 | Bacterial strains, plasmids, and primers used in this study.**

| Strain | Description | Source |
|---|---|---|
| ***P. aeruginosa*** | | |
| PAO-SC4 | AHL synthase-null mutant; PAO1 with unmarked deletions of *lasI* and *rhl* | (Wellington and Greenberg, 2019) |
| PAO-SC4 pPROBE-$P_{rsaL}$ | LasR activity reporter; PAO-SC4 with pPROBE-$P_{rsaL}$ | (Wellington and Greenberg, 2019) |
| PAO1Δ*rhlI* | PAO1 with unmarked deletion of *rhlI* | (Wang et al., 2015) |
| ***E. coli*** | | |
| NEB 5alpha | *fhuA2 Δ(argF-lacZ)U169 phoA glnV44 Φ80 Δ(lacZ)M15 gyrA96 recA1 relA1 endA1 thi-1 hsdR17* | New England Biolabs |
| S17-1 | *recA pro hsdR RP4-2Tc::Mu-Km::Tn7* | (Simon et al., 1983) |
| 5α pJNL pPROBE-$P_{rsaL}$ | LasR-*gfp* reporter; 5α harboring pJNL and pPROBE-$P_{rsaL}$ | (Wellington and Greenberg, 2019) |

| Plasmid | Description | Source |
|---|---|---|
| pJNL | Arabinose-inducible *lasR* expression vector derived from pJN105L(Lee et al., 2006), $Ap^R$ | (Wellington and Greenberg, 2019) |
| pPROBE-$P_{rsaL}$ | *gfp* reporter of LasR activity; pPROBE-GT(Miller et al., 2000) with the *rsaL* promoter extending from -290 to +103, $Gm^R$ | (Wellington and Greenberg, 2019) |
| pJN-lasI | Arabinose-inducible *lasI* expression vector derived from pJN105, $Ap^R$ | This study |
| pJN-empty | Arabinose-inducible expression vector with no gene inserted, ApR | (Wellington and Greenberg, 2019) |
| pJN-RBSlasI | pJN-lasI with native *lasI* RBS | This study |
| pJN-RBSmupI | Arabinose-inducible *mupI* expression vector | This study |
| pEXG2 | Allelic exchange vector with pBR origin, sacB, $Gm^R$ | (Rietsch et al., 2005) |
| pEXG2-lasR | pEXG2 containing *lasR* gene and 500 bp up- and down-stream | [a] |

[a] This plasmid was a gift from M. Kostylev and E.P. Greenberg, constructed using previously published methods (Kostylev et al., 2015).

| Primer Name | Sequence | Description |
|---|---|---|
| lasI-pJN-F | TTGGGCTAGCATGATCGTACAAATTGG | Amplifies *lasI* adding homology to pJN105. *lasI* sequence underlined |
| lasI-pJN-R | TTGGAGCTCCTCATGAAACCGCCAGTC | Amplifies *lasI* adding homology to pJN105, including SacI site. *lasI* sequence underlined. |
| RBS-lasI-F | TTGGGCTAGCAAGGAGGAAGTGAAGATGATCGTACAAATTG | Amplifies *lasI*, including RBS, adding NheI site. *lasI* sequence underlined. |
| mupI-F | TTGGGCTAGCAAGGAGGAAGCCAGCATGAAATATCTAATAG | Amplifies *mupI*, adding RBS and NheI site. *mupI* sequence underlined. |
| mupI-R | TTGGAGCTCTCAAATAGCATTGACTGCGTCC | Amplifies *mupI*, adding SacI site. *mupI* sequence underlined. |
| mupI-pJN-F | TCCATACCCGTTTTTTTGGGCTAGCAAGGAGGAAGCCAG | Amplifies *mupI* PCR product produced by mupI-F & mupI-R, adding additional homology to pJN105 |
| mupI-pJN-R | CTCACTATAGGGCGAATTGGAGCTCCTCAAATAGCATTG | Amplifies *mupI* PCR product produced by mupI-F & mupI-R, adding additional homology to pJN105. *mupI* sequence underlined. |

*Site-directed mutagenesis primer sequences are available upon request.*

**a**

| LasI amino acid position | LasR amino acid position | GREMLIN score |
|---|---|---|
| 142 | 127 | 0.105 |
| 145 | 61 | 0.096 |
| 157 | 129 | 0.096 |
| 102 | 61 | 0.092 |
| 140 | 130 | 0.091 |
| 103 | 91 | 0.090 |
| 152 | 36 | 0.090 |
| 144 | 76 | 0.090 |
| 142 | 64 | 0.088 |
| 144 | 129 | 0.086 |
| 20 | 237 | 0.086 |
| 69 | 127 | 0.086 |
| 145 | 36 | 0.084 |
| 46 | 125 | 0.083 |
| 99 | 50 | 0.083 |
| 181 | 148 | 0.083 |
| 159 | 178 | 0.082 |
| 75 | 57 | 0.081 |
| 125 | 75 | 0.081 |
| 69 | 36 | 0.080 |
| 102 | 38 | 0.079 |
| 138 | 66 | 0.078 |
| 16 | 228 | 0.078 |
| 136 | 197 | 0.077 |
| 148 | 76 | 0.077 |
| 155 | 70 | 0.076 |
| 145 | 213 | 0.076 |
| 152 | 38 | 0.076 |
| 41 | 183 | 0.075 |
| 16 | 207 | 0.074 |

**b**

| LuxI amino acid position | LuxR amino acid position | GREMLIN score |
|---|---|---|
| 143 | 137 | 0.109 |
| 146 | 44 | 0.103 |
| 102 | 46 | 0.100 |
| 57 | 147 | 0.091 |
| 103 | 99 | 0.091 |
| 146 | 69 | 0.090 |
| 126 | 83 | 0.087 |
| 114 | 206 | 0.087 |
| 145 | 84 | 0.086 |
| 54 | 126 | 0.084 |
| 171 | 211 | 0.084 |
| 102 | 69 | 0.080 |
| 158 | 139 | 0.080 |
| 69 | 137 | 0.079 |
| 126 | 43 | 0.079 |
| 141 | 140 | 0.078 |
| 67 | 162 | 0.078 |
| 75 | 65 | 0.078 |
| 143 | 72 | 0.076 |
| 145 | 139 | 0.076 |
| 156 | 78 | 0.076 |
| 139 | 74 | 0.075 |
| 185 | 178 | 0.075 |
| 146 | 221 | 0.075 |
| 115 | 148 | 0.074 |
| 115 | 26 | 0.074 |
| 17 | 163 | 0.074 |
| 67 | 182 | 0.074 |
| 106 | 224 | 0.074 |
| 84 | 127 | 0.073 |

**c**

```
LasI  59 DTPEAQVFGCWR ILDTTGPYML KNTFPELLHG KEAPCSPHIW ELSRFAINSG 110
LuxI  61 DT--ENVSGCWR LLPTTGDYML KSVFPELLGQ QSAPKDPNIV ELSRFAVGKN 110

LasI 111 QKGS-LGFSD CTLEAMRALA RYSLQNDIQT LVTVITVGVE KMMIRAGLDV 159
LuxI 111 SSKINNSASE ITMKLFEAIY KHAVSQGITE YVTVTSTAIE RFLKRIKVPC 160
```

**d**

```
LasR     1 MAL-----VD GFLELE---R SSGKLEWSAI LQKMASDLGF SKILFGLLPK 42
LuxR     1 MGMKDINADD TYRIINKIKA CRSNNDINQC LSDMTKMVHC EYYLLAIIYP 50

LasR    43 DSQDYENAFI VGNYPAAWRE HYDRAGYARV DPTVSHCTQS VLPIFWEPSI 92
LuxR    51 HSMVKSDISI LDNYPKKWRQ YYDDANLIKY DPIVDYSNSN HSPINWNIFE 100

LasR    93 Y--QTRKQHE FFEEASAAGL VYGLTMPLHG ARGELGALSL SVEAENRAEA 140
LuxR   101 NNAVNKKSPN VIKEAKTSGL ITGFSFPIHT ANNGFGMLSF AHSEKDNYID 150
```

**Supplementary Fig. 1 | Top-scoring GREMLIN residues.** Top-scoring covarying amino acid residues identified in **a)** *P. aeruginosa* PAO1 LasI/R or **b)** *V. fischeri* MJ11 LuxI/R along with the GREMLIN score (with APC) for each pair. Residues that appear on both lists are shown in boxes on the sequence alignments **c)** of LasI and LuxI and **d)** of LasR and LuxR. Only relevant portions of the protein sequences are shown.

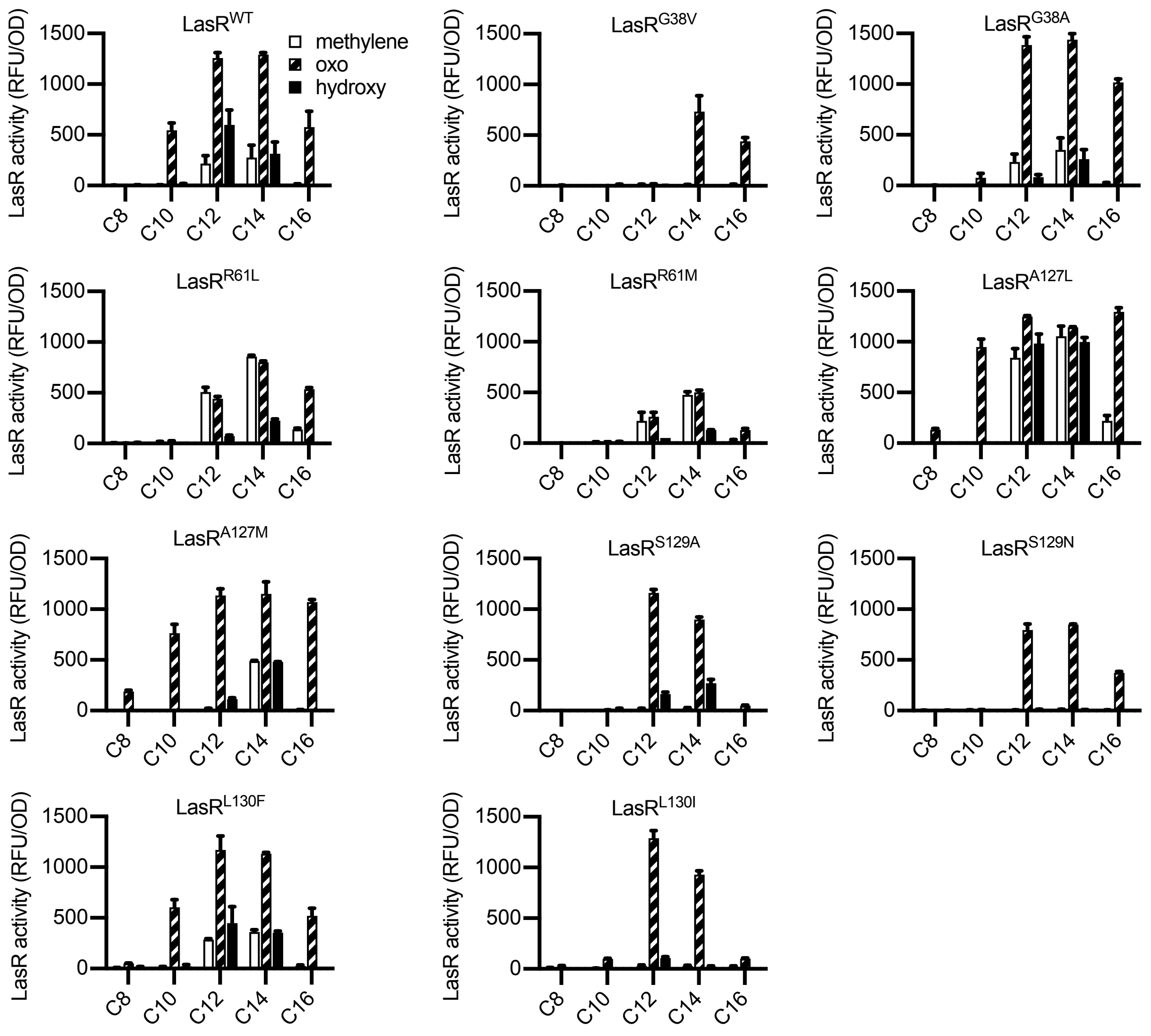


**Supplementary Fig. 2 | Activity of LasR mutants in *E. coli*.** LasR activity in response to a panel of AHL signals (100 nM; chain length indicated on horizontal axis, C3 modification indicated by shading) is reported as relative fluorescence units (RFU) normalized by optical density at 600 nm (OD). Wild-type (WT) or *lasR* with the indicated amino acid substitution were expressed from pJNL in *E. coli* NEB 5alpha harboring pPROBE-$P_{rsaL}$. Signals with 4 or 6 carbons in the acyl chain did not activate any of the LasR variants and are not shown. The following LasR mutants had little or no activity in response to 100 nM AHLs: R61V, R61Q, S129T, G38L. Data are the mean and standard deviation of two biological replicates and are representative of three (mutants) or ten (WT) independent experiments.

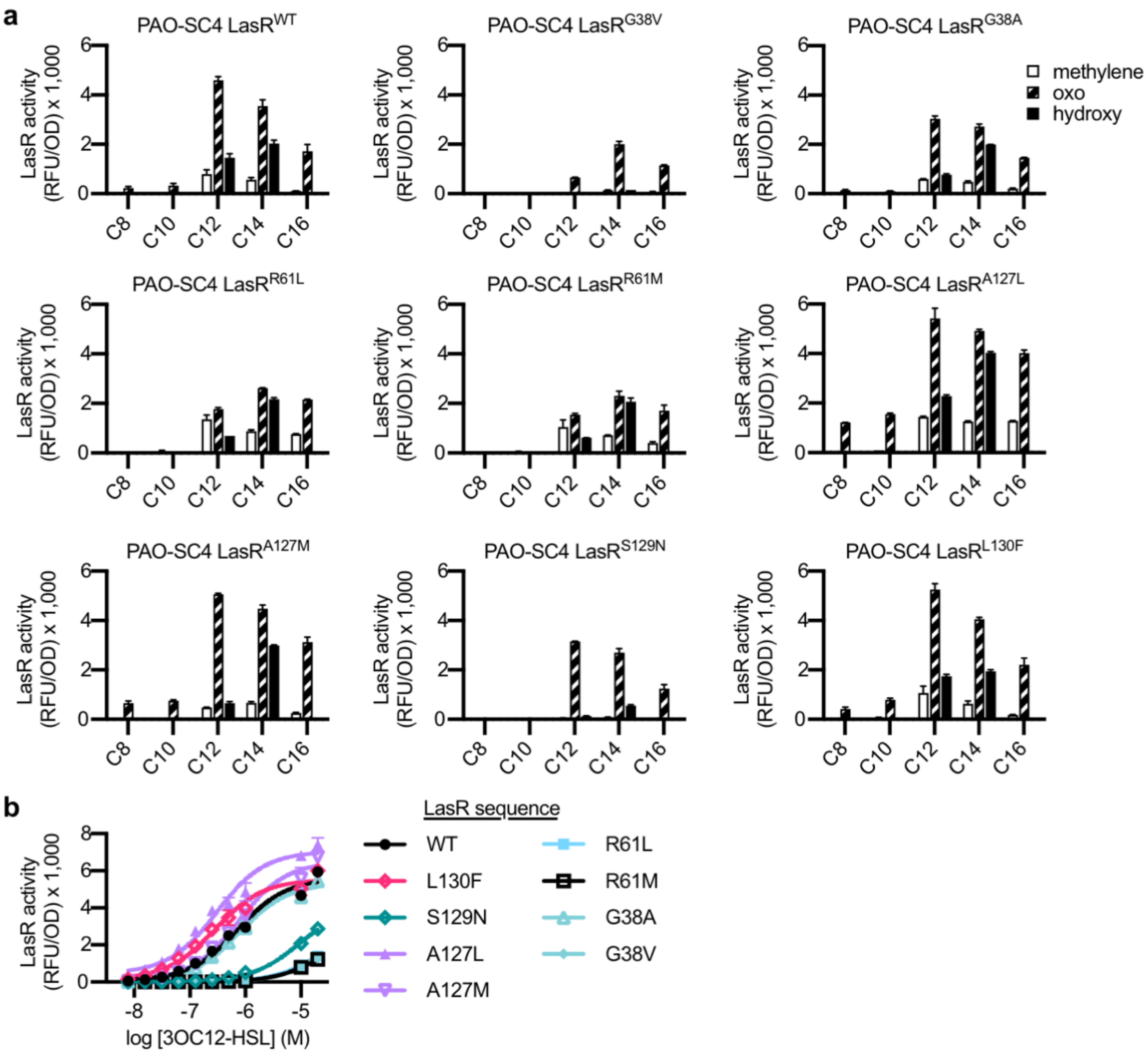


**Supplementary Fig. 3 | Activity of *P. aeruginosa* LasR mutants.** The activity of unmarked, chromosomal *lasR* mutants of *P. aeruginosa* PAO-SC4 containing pPROBE-$P_{rsaL}$ measured as GFP fluorescence (relative fluorescence units normalized to optical density at 600 nm, RFU/OD). **a)** Activity of each LasR mutant or wild-type (WT) in response to a panel of 19 AHLs at 20 μM. AHLs with 4 or 6 carbons in the acyl chain did not activate any of the LasR variants and are not shown. Data are the mean and standard deviation of two biological replicates and are representative of three independent experiments. **b)** Activity of each LasR mutant or WT in response to 3OC12-HSL. Data are the mean and standard deviation of three biological replicates and are representative of three (mutants) or four (WT) independent experiments.

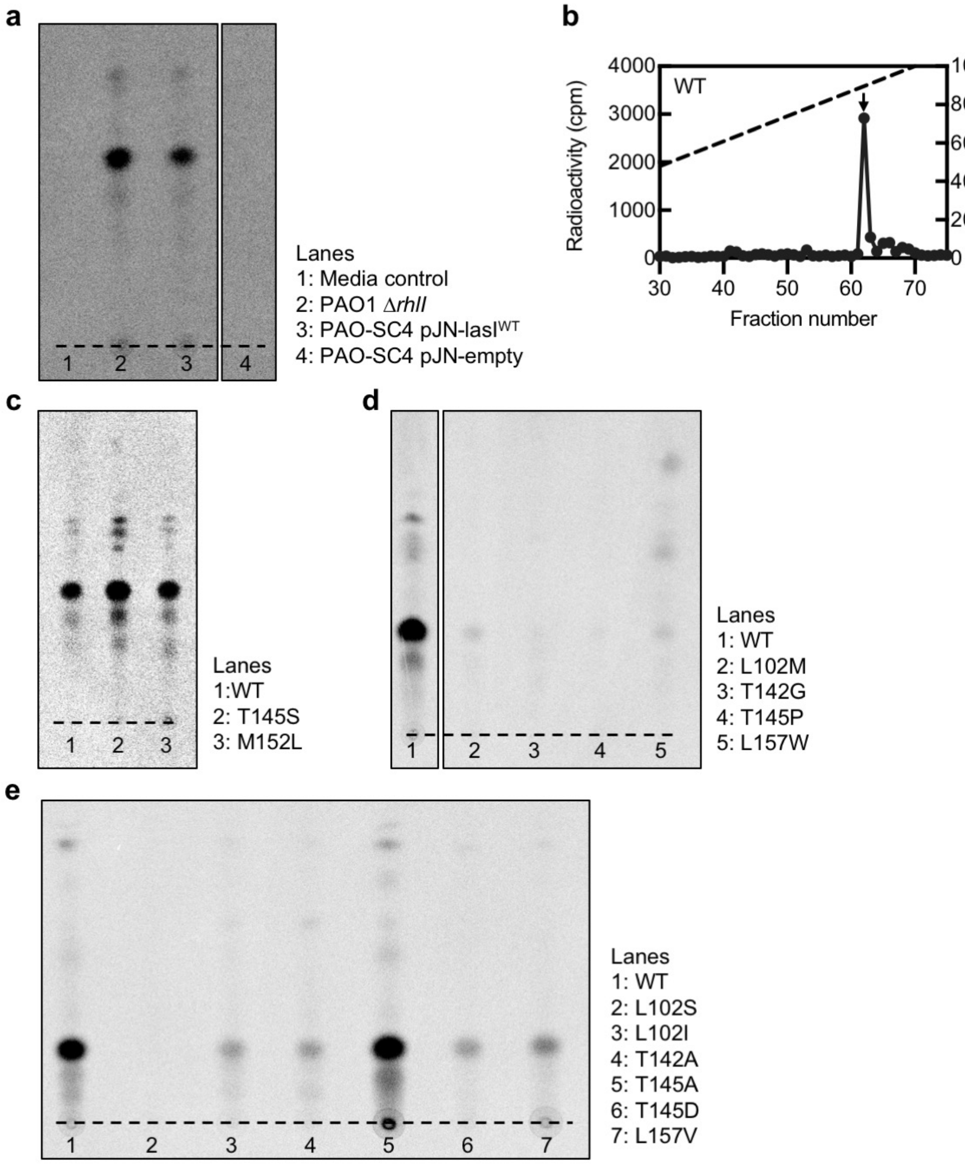


**Supplementary Fig. 4 | Radiotracer assays of LasI activity. a)** Thin layer chromatography (TLC) analysis of $^{14}$C-AHLs extracted from *P. aeruginosa* PAO1ΔrhlI or from *P. aeruginosa* PAO-SC4 harboring pJN-lasI$^{WT}$ or pJN-empty or from a media control in which no cells were present. The dashed lines indicates the origin. Results are representative of two independent experiments. **b)** Extracts from *P. aeruginosa* PAO-SC4 pJN-lasI$^{WT}$ shown in panel **a** were also analyzed by high performance liquid chromatography (HPLC). The horizontal axis denotes the fraction number; fractions 1-29 are not shown. The dashed line indicates the methanol gradient, plotted on the right vertical axis. The counts per minute (cpm) of radioactivity in each fraction are plotted on the left vertical axis. The arrow indicates the fraction in which 3OC12-HSL elutes. **c-e)** TLC analysis of $^{14}$C-AHLs extracted from *P. aeruginosa* PAO-SC4 pJN-lasI WT or with the indicated amino acid substitution.

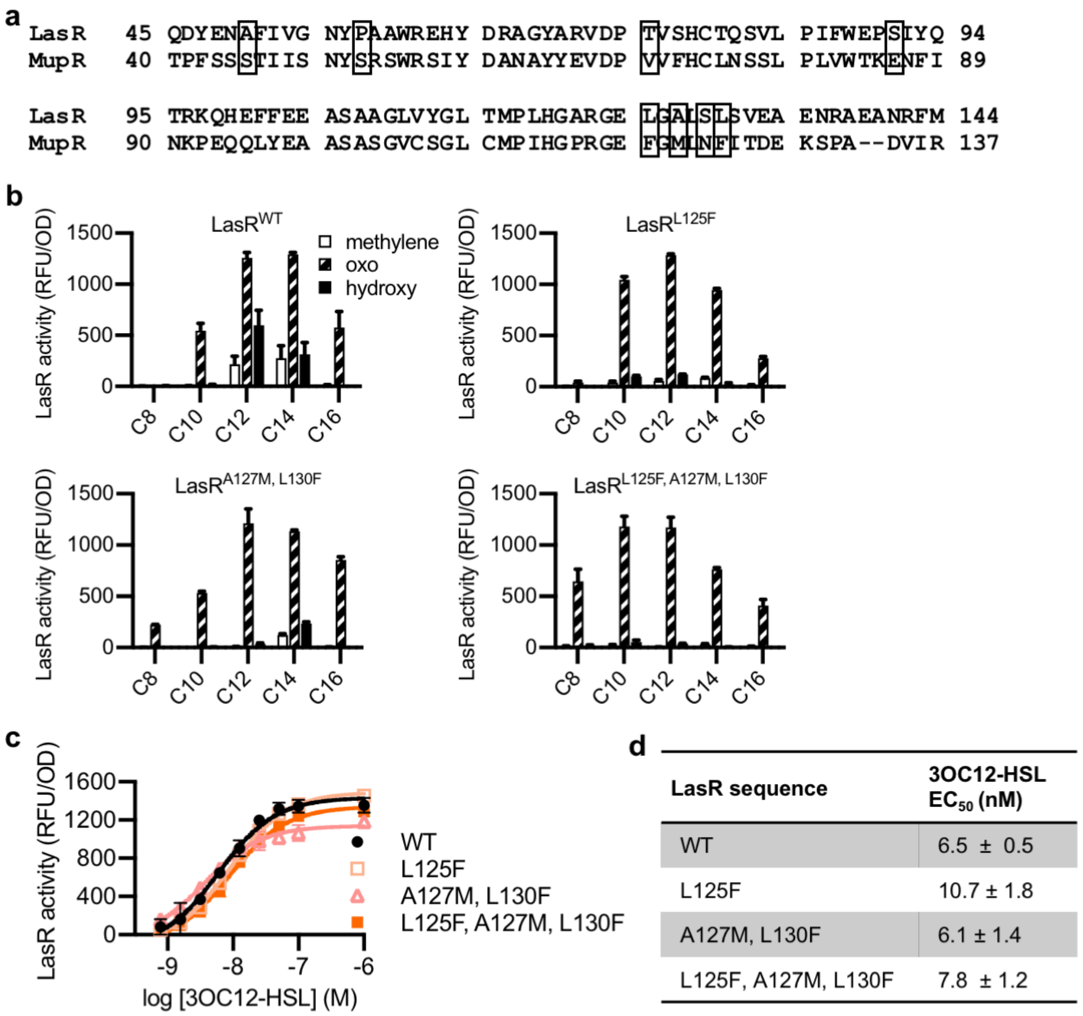


| LasR sequence | 3OC12-HSL $EC_{50}$ (nM) |
|---|---|
| WT | 6.5 ± 0.5 |
| L125F | 10.7 ± 1.8 |
| A127M, L130F | 6.1 ± 1.4 |
| L125F, A127M, L130F | 7.8 ± 1.2 |

**Supplementary Fig. 5 | "MupR-like" LasR mutant activity. a)** Sequence alignment of LasR and MupR. Boxes indicate covarying residues with a GREMLIN score (with APC) >0.08 that differ between the two proteins. **b)** Activity of LasR (wild-type, WT, or mutated) measured in *E. coli* pJNL pPROBE-PrsaL in response to a panel of 19 AHL signals at 100 nM. AHLs with 4 or 6 carbons in the acyl chain did not activate any of the LasR variants and are not shown. LasR activity is reported as relative fluorescence units (RFU) normalized to optical density at 600 nm (OD). Data are the mean and standard deviation of two biological replicates and are representative of three (mutants) or ten (wild-type) independent experiments. **c)** Activity of wild-type (WT) or mutant LasR measured in *E. coli* pJNL pPROBE-PrsaL in response to 3OC12-HSL. Data are the mean and standard deviation of three biological replicates and are representative of three independent experiments. **d)** Concentration of half maximal activity ($EC_{50}$) of 3OC12-HSL for LasR, calculated from data shown in panel **c**. Data are mean and SEM.

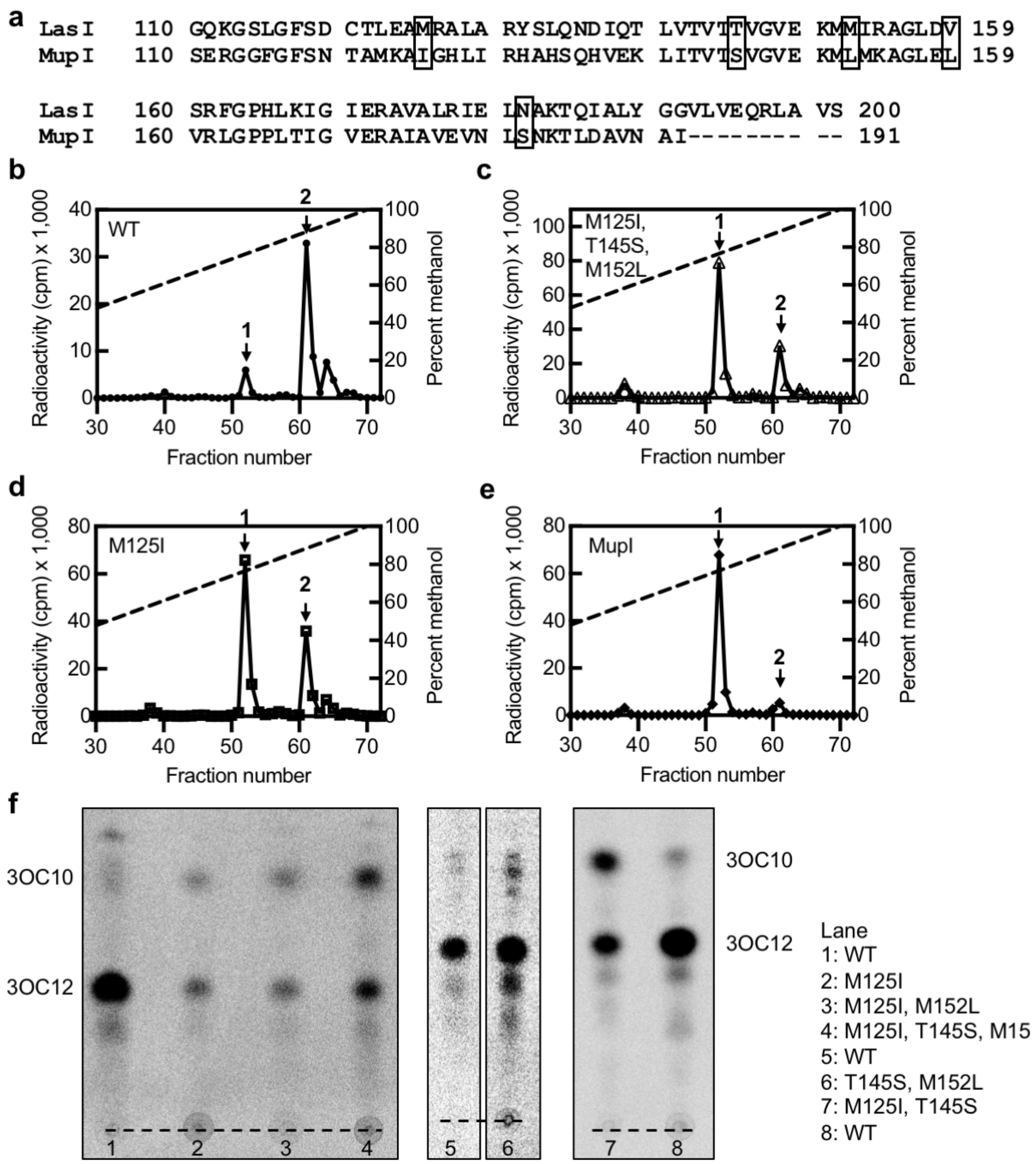


**Supplementary Fig. 6 | "MupI-like" LasI mutant activity. a)** Sequence alignment of LasI and MupI. Boxes indicate covarying residues with a GREMLIN score (with APC) >0.08 that differ between the two proteins. HPLC analysis of $^{14}$C-AHLs extracted from *P. aeruginosa* PAO-SC4 harboring **b)** pJN-RBSlasI$^{\mathrm{WT}}$ **c)** pJN-RBSlasI$^{\mathrm{M125I,T145S,M152L}}$, **d)** pJN-RBSlasI$^{\mathrm{M125I}}$, or **e)** pJN-RBSmupI. The horizonal axis denotes the fraction number; fractions 1-29 are not shown. The left vertical axis indicates the counts per minute (cpm) of radioactivity in each fraction. Arrow 1 indicates the fraction at which 3OC10-HSL elutes and arrow 2 indicates the fraction at which 3OC12-HSL elutes. Data are representative of two (mutants) or three (wild-type, WT) independent experiments. **f)** TLC analysis of $^{14}$C-AHLs extracted from *P. aeruginosa* PAO-SC4 pJN-lasI wild-type (WT) or with the indicated mutations. The identities of the two major spots on each TLC were deduced from HPLC analysis of the same extracts.
